# Differentiation smelling footprints of the Chagas disease vector using an electronic nose based on artificial intelligence algorithms

**DOI:** 10.1101/2024.02.13.579939

**Authors:** Luisa F. Ruiz-Jiménez, Daniel A. Sierra, Homero Ortega B, Bladimiro Rincon-Orozco, Jonny E. Duque

## Abstract

The present study aims to present the design of an electronic nose capable of learning and differentiating semiochemical signals emitted by insects usable to identify species that transmit Chagas disease. The proposed device used different non-specific resistor gas sensors integrated into a system of artificial intelligence models. To validate the nose, we used eight insect species of the Triatominae subfamily and one population that was a natural carrier of the parasite *Trypanosoma cruzi*. Also, the discriminatory capacity of distant species was tested with other insects like *Aedes aegypti* (arbovirus vector) and *Sitophilus oryzae* (stored grains plague). As a result, the electronic nose was able to differentiate up to gender level with an accuracy of 89.64% and to differentiate *Rhodnius pallensces* naturally infected with *T. cruzi* with less than 1% of error in classification. These results show that our designed device can detect particular smelling footprints, and one electronic nose like that could be a tool to discriminate against insects in the future.

## Introduction

Every year, more than one million deaths worldwide are recorded due to contagion with vector-borne diseases, representing approximately 17% of widespread infectious diseases in humans. It is highlighted that the growth of this threat is linked to world globalization, as has happened with such diseases as chikungunya, Ebola, influenza H1N1, or inclusive COVID-19 (Carcavallo and Rabinovich 1985; Morens and Fauci 2013; Vesga et al. 2022). The Chagas disease is an excellent example of the huge problem caused by insects’ hematophagous. People can be infected with the intracellular flagellate parasite *Trypanosoma cruzi* (Kinetopastisida: Trypanosomatidae) that can be transmitted by insect species belonging to the Triatominae subfamily (Hemiptera: Reduviidae) (Guarneri and Lorenzo 2021). These insects are popularly called kissing bugs, and their diet is on blood suction. However, the infection comes just from the feces of the vector infected with trypomastigotes, which are left after biting. Feces spread through the wounded skin after entering the organism due to the intense rubbing that follows the pruritus of the wound. In addition to vector transmission, there are other routes of contagion, such as contamination by food or by mouth, blood transfusions, organ transplants, and laboratory accidents (Moncayo and Silveira 2017).

This sickness is present from southern California – the USA to central regions of Argentina and Chile, affecting approximately 21 Latin American countries in the Neotropical area. About 6 million to 7 million people worldwide are estimated to be infected with *T. cruzi*. Because of human migration, non-native infection cases have been found in non-endemic places such as the United States, Canada, Japan, and some European and Western countries (Requena-Méndez et al. 2015; Moncayo and Silveira 2017). In Latin America, the three countries with the most significant were Argentina with 1.505.235 cases, Brazil with 1.156.821, and Mexico with 876.458 cases, respectively. On the other hand, in the Andean Region, about 958.453 people were infected, 45.7% of them (437,960) were living in Colombia, where the disease turned out to be endemic. In Colombia, the Chagas parasite transmission has been reported in more than 15 states in the eastern part of the country. The state of Santander, where the present study was held, is one of the most endemic in the country (Reyes et al. 2017). This disease can be asymptomatic for years, leaving severe sequelae in organs like the colon and heart. Chagas disease has two stages or clinical phases: an acute phase and a chronic phase. It is important to highlight that many people (70 to 80% of those infected) are asymptomatic throughout their lives. However, 20 to 30% of those affected by this disease evolve into chronic symptomatic pictures associated with heart, digestive tract, and nervous system damage. Chagas disease also affects the central nervous system, the enteric nervous system, and the digestive system (Malone et al. 2021).

As a fundamental pillar to designing prevention and control programs to control vector-borne diseases, it is necessary to study the taxonomic identification, habitat, bionomy, eating habits, dispersion of capacity, and life cycles of these insects, among other biological characteristics (Guarneri and Lorenzo 2021). For example, even identifying species is challenging for experts since more than 3,000 species of vectors are distributed globally. This is why it is an exclusive work for entomologists, each with deep knowledge of a particular species. Here is where the contribution of artificial intelligence becomes relevant to facilitate species recognition to non-specialists in these areas. For this reason, an automated system able to discriminate the taxonomy of insects through the detection of chemical compounds issued to the environment will be a good tool for pest management control decisions. Today, the fourth industrial revolution allows us to establish a prospect where advances in specialized humanoid robotics technology make feasible solutions to all previously discussed challenges. Developing new intelligent materials and artificial intelligence (machine learning) associated with the Internet of Things (IoT) may contribute to a new revolution in medical entomology, especially in recognizing vector insects.

As previously mentioned, it is of great interest to medical entomology to know the aspects related to the behaviors, orientation, and assertive identification of triatomines (Lopes et al. 2020) ^10^. In several research works, evidence of chemical sensory signals emanated by the insects as a means of communication has been found (Tabata 2018), so they can locate hosts, shelters, alert signals, and sexual partners, among other beneficial behaviors for the individual and the species. The presence of volatile chemical compounds secreted by kissing bugs has been related to the scent of the metasternal and Brindley glands (Walker et al. 2016; Anton et al. 2017; Guarneri and Lorenzo 2021). However, independently of the structure responsible for producing and secreting info-chemicals (semiochemicals), there is a distinctive odor, rancid and pronounced, generated by individuals and colonies of different species of triatomines (Schuh and Slater 1995). An electronic nose is a device composed of gas sensors that respond with electrical signals in the presence of certain specific chemical compounds according to their configuration (Gardner and Bartlett 1994). The generated electrical response is then fed to a classifier based on machine learning algorithms for pattern recognition (Kim et al. 2021). The expectative with these devices is that they allow the joint analysis of the molecules between vectors and other insect non vectors, creating a unique smelling footprint for each factor studied.

Currently, there are numerous applications of the electronic nose concept (Zhang et al. 2018), where their primary use ranges go from the study and identification of diseases (Cumeras and Correig 2018), agro-food industry (Denizli et al. 2021) work with cosmetics (Abedi et al. 2018), oils (Buratti et al. 2018), wines, and coffee, control of environmental contaminants (Vedal et al. 2017), process monitoring, detection of explosives, toxic substances and drugs (Yinon 1999). In short, electronic noses are inspired by the ability of bloodhounds to identify objects under challenging situations thanks to their sense of smell (Vorobioff et al. 2018). In consequence, to replicate the functioning of a biological olfactory system from the acquisition of volatiles to the identification of odors, critical limitations (Rajamäki et al. 2006) have to be addressed, mainly associated with the technological advance of the sensors capable of detecting volatiles at low concentrations or specifically distinguish an only odor of a mixture of odorants (Gardner and Bartlett 2013).

As a preamble to future studies in the automatic taxonomic detection of vector disease insects with artificial intelligence and to support the convergence of new technologies associated with public health, this work aims to propose an electronic olfactory device, called an electronic nose, that can learn to discriminate between the odor produced by colonies of different species of triatomines and the odor produced by species of other taxonomic orders.

## Materials and Methods

The research was developed in the Universidad Industrial de Santander insectarium in Bucaramanga, Colombia. The electronic nose construction materials are described below as an IoT device that detects triatomines. The methodology implied data collection, experiment, pattern recognition, and statistical analysis.

### Principle of Operation And Model of the Electronic Nose

Like all electronic noses, the proposed device is based on three types of interactions: chemistry, electronics, and logical support (software and learning algorithms) (Moreno et al. 2009; Vorobioff et al. 2018), which are described at the level of physical components and programs-routines (in the block diagram) as can be seen in figure S4.

The chemical stimulus begins with the oxidation-reduction reaction between the sample to be studied and the resistive gas sensors. For this, the volatile study is conditioned (chemical ecology produced by the triatomines and other odors), which is redirected to a combined matrix of materials sensitive to different chemical compounds. The matrix sensors present measurable variations in their conductivity levels, generating analog signals processed at the hardware and software levels during electronic interactions.

The prototype built has a matrix of 16 semiconductor sensors, which work with a film composed of oxide-metal crystals type n, in this case, tin dioxide (SnO2). When a metal oxide crystal such as SnO2 increases its temperature, the oxygen in the air is absorbed on the surface of the crystal with a negative charge. Subsequently, the donor electrons on the surface of the crystal are transferred to the absorbed oxygen and have a higher positive charge, which means an increase in the potential barrier as well as the electrical resistance of the material. In the presence of a deoxidizing gas, the surface density of negatively charged oxygen decreases, as does the potential barrier, and therefore the conductance of the material increases. These variations respond to the concentration of the gas in measurement (Göpel et al. 1989).

The electronic interactions were based on digitizing the analog response produced by each of the 16 sensors as signal conditioning. This phase is produced by the ADC (Analog Digital Converter) development board, which uses serial data sending (SPI) protocol to communicate with the Intel Galileo Generation 1 development card. This card connects to the Internet through DHCP (Dynamic Host Configuration Protocol) via an Ethernet connector; its logic of sending and receiving data is confined to a simple local routine written in Python language that allows the remote control of the electronic nose. The process begins with the reception of the digital signals of the 16 channels corresponding to the response of the 16-sensor array, which is then housed in the database (Electronic Nose Data Base) in the form of a data matrix whose rows correspond to the unit of sampled time acquired and the columns to each one of the channels.

The first computational step is the application of a Butterworth digital low pass filter with a cutoff frequency Wn = 0.2 applied to each signal. Subsequently, for the electronic nose in learning mode, the extraction of characteristics is performed, followed by the selection of relevant characteristics, classification, training, and finally, the validation that allows deducing the performance measures of the classifier. The parameters used in the learning stage are loaded from the database, and the classifier is evaluated to obtain the identification result for the smell recognition stage.

The electronic nose needs three minimum periods to measure a sample: a time of concentration of molecules and stabilization (tc) to establish the value of basal resistance of the sensors and, at the same time, concentrate as much as possible the volatiles in a specific volume; sensing time (ts) where the sensor data is acquired and stored in the database; and finally, a cleaning time (tl) is determined to remove the chemical compounds measured and oxidize-reduce the material of the sensors, bringing them to their resting state.

### Electronic Nose Hardware

They are considering that the application of the electronic nose to identify triatomines benefits from the IoT mobile terminal concept in terms of communication. A user profile is created for the developers and another for users, with their respective database addresses and web access. The hardware of the designed electronic nose was based on similar models proposed in previous engineering works, combined with experiences of device migration towards IoT platforms (José de Jesús et al. 2016)74,75. The electronic nose consists of a sensing module, an energization module, and an acquisition-communication module, as shown in Figure S4.

### Biologic Material

Colonies belonging to the CINTROP research center’s insectarium were used in the experiments, and the infected biological material was collected in parasite transmission areas (Jagua de Ibirico, Cesar, Colombia). For each triatomine species, different colonies are found in wide-mouth plastic bottles, as shown in Figure S3 (A to F). The elements available are a cardboard base that efficiently collects the biological waste generated by individuals, such as feces, exuviae, or corpses, and fan-shaped cardboard with hexagonal or circular holes that allow them to transit through the container. This fan shape is due to the behavioral characteristic of defense since they mainly seek places such as corners, edges, or cracks. Finally, to keep the individuals inside the container, light fabric with an open structure in the shape of a net (’tul veil’) of white color (to generate contrast and easily observe the triatomines) and with a maximum distance between the weft threads of 1.5mm, the fabric is secured with thick rubber and masking tape around, to prevent the escape of insects. During the procedure, law 84 of 1989 of chapter VI, articles 23 (literal a-b-c) and 24 of the National Statute for the Protection of Animals, which regulates the use of live animals in research experiments compiled. Complying with the laboratory animal handling procedures and under the approval of the Ethics Committee “Comité de ética en investigación científica de la Universidad Industrial de Santander CEINCI”: Scientific Research Ethics Committee of the Industrial University of Santander (acronym in Spanish CEINCI); Minutes No. 08, May 11, 2018 - see supplementary document of approval S5). Furthermore, the study was carried out in compliance with the Arrive guidelines.

In the insectary, four types of containers are handled, as shown in Figure S3G. Type A corresponds to the smallest volume, this being 1.14 L (11 cm in diameter by 12 cm in height), followed by type B of 2.31 L with dimensions of 14.5cm in diameter and 14 cm in height, type C corresponds to 3.63 L in volume, 17 cm in diameter and 16 in height. Finally, type D has 5.65 L of volume and is 20 cm in diameter by 18 cm in height.

### Laboratory Conditions

All the experimentation carried out with the electronic nose was performed at night, between 5:00 p.m. and 7:00 p.m. (the following day), to cover most of the activity produced by the triatomines (Lazzari 1992; Mosqueiro and Huerta 2014)76,59,58, since at night, the behavior of searching for food, couple, shelter among others increases. During these hours, the laboratory remained alone with the presence of the person who applied the experiments, presenting a temperature variation between 22.7 ° C - 23.8 °C and relative humidity levels between 72.5% - 77.7%. The laboratory’s temperature and humidity levels were measured using an Extech Instruments brand thermo-hygrometer and tagged 42280. Its location was always as close as possible to the sensors.

PCR was performed as described previously (Fraga et al. 2014) briefly, extracted DNA from insect intestines and was amplified by PCR using the following primers: f: GCAACCAGATTGTCATCACGAACG and r: GTCCTGCGMCTCGTACTTGGCA and this conditions 0.8 uM primers, 2.5 mM MgCl2, one cycle 95 °C for 5 min, 35 cycles 94 °C 40 s/ 66 °C 1 min/ 72 °C and 1 min 1 cycle at 72 °C 10 min after that amplified PCR products were detected in a 2% agarose gel stained with Syber green.

### Database Creation

The experiment’s design was based on the data acquisition of 3 replicates and three repetitions of each associated factor. A random list with a normal distribution of the list of units (number of replicates of trials corresponding to treatment factors of the positive class - with triatomines) is established. The repetitions correspond to immediate consecutive acquired measurements, and the replicas were made on different days. The database has 684 files (total number of tests) typed in plain text (.csv). Each is a data matrix of 16 columns (the 16 channels of each sensor) and approximately 1000 rows, corresponding to an acquisition time of 10 minutes: with a sampling frequency of 1.7 Hz. All data generated and analyzed during this study are included in this published article as a supplementary file S6.

### Statistical Analysis and Data Processing (Electronic Nose Logical Support)

Electronic noses are commonly associated with data processing based on pattern recognition. This consists of extracting information from the data to establish properties among the analyzed objects, thus inducing sets with the same characteristics (Rodriguez-Lujan et al. 2014; Mosqueiro and Huerta 2014). That is, it gives the nose the ability to detect, quantify, and identify particular smells. The data processing of the prototype is programmed in Python language and consists of the stages of extraction of principal characteristics, selection of the most relevant characteristics, classification of data, training of the machine, and finally, validation of learning.

The feature extraction stage includes compensation of the sensor’s drift caused by temperature and humidity variability in the environment. The conductance response curve (inverse of the resistance) varies over time as the material of the sensor is oxidized/reduced. The parameters studied for the extraction of characteristics were the average of the first 200 samples as the initial reference conductance; the final conductance value; the maximum conductance value and the time in which said conductance value was presented; the arithmetic mean of the initial response and reference conductance vector; and finally, the standard deviation of the initial response and reference conductance vector.

Different characteristics of the signals do not imply that they represent most of the classificatory information. Therefore, the feature selection stage reduces the number of variables, eliminating redundant ones and those that do not provide additional information for separability. The principal component analysis (PCA) technique is used, which reduces the dimensionality to a set of data depending on the variance value from the covariance matrix of the signals.

Finally, the learning algorithms based on support vector machines (SVM) are applied, which make up the family of the most powerful and popular techniques of machine learning thanks to their robustness. The algorithms were developed based on the ‘scikit-learn v0.19.1’ library of Python (Pedregosa et al. 2011; Vembu et al. 2012), where the programming of vector support machines with linear kernel, polynomial, radial, and an exception in the linear kernel *one-vs-rest* called ‘LinearSVC’ are housed.

In the same way, as in the case of the classification and training stage, the ‘scikit-learn v0.19.1’ library of Python (Pedregosa et al. 2011; Vembu et al. 2012) allows cross-validation of the trained machines. This validation is iterated 500 times to estimate, in each case, the highest error (accuracy) that machines can commit. The sample size to be evaluated corresponds to 30% of the full sample size. At the end of the process, the vector support machine is chosen with the kernel with the lowest prediction error.

## Results

The main results of experiments carried out with the electronic nose built as an IoT device are presented to evaluate the ability of the nose to differentiate triatomine odors from interfering odors from other kinds of insects. The learning system based on pattern recognition is also presented in conjunction with the different tests needed to reach, in each case, the performance levels needed for the successful discrimination and classification of the odors of interest. In particular, the kissing bugs, mosquitoes, and rice weevils used in the tests have been maintained under laboratory conditions to study their odors, except for the naturally infected Triatominae colony obtained from areas of parasite circulation.

### Operating Principle of the Electronic Nose – Testing with Chemical Solvents

The main element of a nose is a set of different odor sensors. An odor sensor is a gas sensor considering the tendency of manufacturing specialized gas sensors, which can respond differently to different gasses. A gas sensor is an electrical conductor whose conductivity is sensitive to particular gas or gasses so that an electrical signal passing through it varies depending on conductivity changes and, consequently, on changes in that gas or gas presence. It is the sensor response or the sensor conductance response. A set of different odor sensors is what, from now on, will be called a sensor matrix. The matrix uses sensors of different characteristics, giving a spectrum of responses specific to a specific odor of interest. But an odor of interest usually corresponds to an object of interest. Using a proper number of different sensors, the spectrum can be seen as a scented print for an object of interest, equivalent to humans’ fingerprints. Like bloodhounds with different nose recognition characteristics but can be trained to do the same work, two sensor matrices with different characteristics can perform the same pattern recognition task due to a training process.

Thus, in several interactions, an electronic nose must detect and treat a set of electrical signals by hardware and software. The prototype used had a matrix of 16 built-in semiconductor sensors, one working with a film composed of n-type metal oxide crystals, in our case, tin dioxide (SnO2). When a metal oxide crystal such as SnO2 increases its temperature, the oxygen in the air is absorbed on the surface of the crystal with a negative charge. Subsequently, the donor electrons on the surface of the crystal are transferred to the absorbed oxygen with a higher positive charge, which increases the potential barrier and the material’s electrical resistance. In the presence of a deoxidizing gas, the density of the negatively charged oxygen surface decreases, as does its potential barrier. Therefore, the conductance of the material increases. These variations respond to the concentration of the gas present.

An electronic nose’s electronic process begins with converting the electric signals to numerical data. It is done by a data communication system that sends the 16 sensor signals to a cloud server by applying modern digital processing methods, including artificial intelligence. Testing the operating principle of the electronic nose is just the first step to be done and means not only observing the proper functioning of the sensors and the system but also getting a characterization of the sensor matrix. It is essential to note that sensors’ characteristics change in time due to wear and tear when used in different conditions. This step must be repeated periodically as part of a synchronization process for the nose. Different solvents in high concentrations (> 90%) are proposed for the test to characterize the sensors represented by their conductance response to those critical solvents. Four different types of actions were chosen due to the polarity, the number of electrons that allow the creation of links with the elements present in the material of each sensor (SnO2), and the volatility factor. This way, the operating status of the electronic nose as an IoT device was mapped, which will help control temporary wear drifts and identify irregular behavior in the sensors’ measurements in time. The chosen compounds correspond to acetonitrile, methanol, acetone, and dimethyl-sulfoxide.

Figure 1 (A to D) shows the chemical structure of each solvent used in the synchronization tests, followed by the conductance responses of the gas sensors whose signals have been normalized and centered. The graphs correspond to the box plot that provides an overview of the symmetry of the response distribution of the sensors, where a particular and different scent print for each chemical compound is identified. The dispersion values represent the changes in amplitude and speed in the response of the sensors. For the case of acetone, the sensor S12 (tagged as TGS826) presented a peak response with a slope high enough to locate said values as outliers; however, 75% of the response of the sensor was located around the zero value, that is, the sample volatilized with the certain speed that the sensor stopped detecting it quickly. In this way, the location of the interquartile range is related to the volatility factor of the compound, as observed in the case of DMSO (dimethyl-sulfoxide), whose response is in the stable state of the sensors is different from zero, that is, to the basal state.

**Fig. 1.**
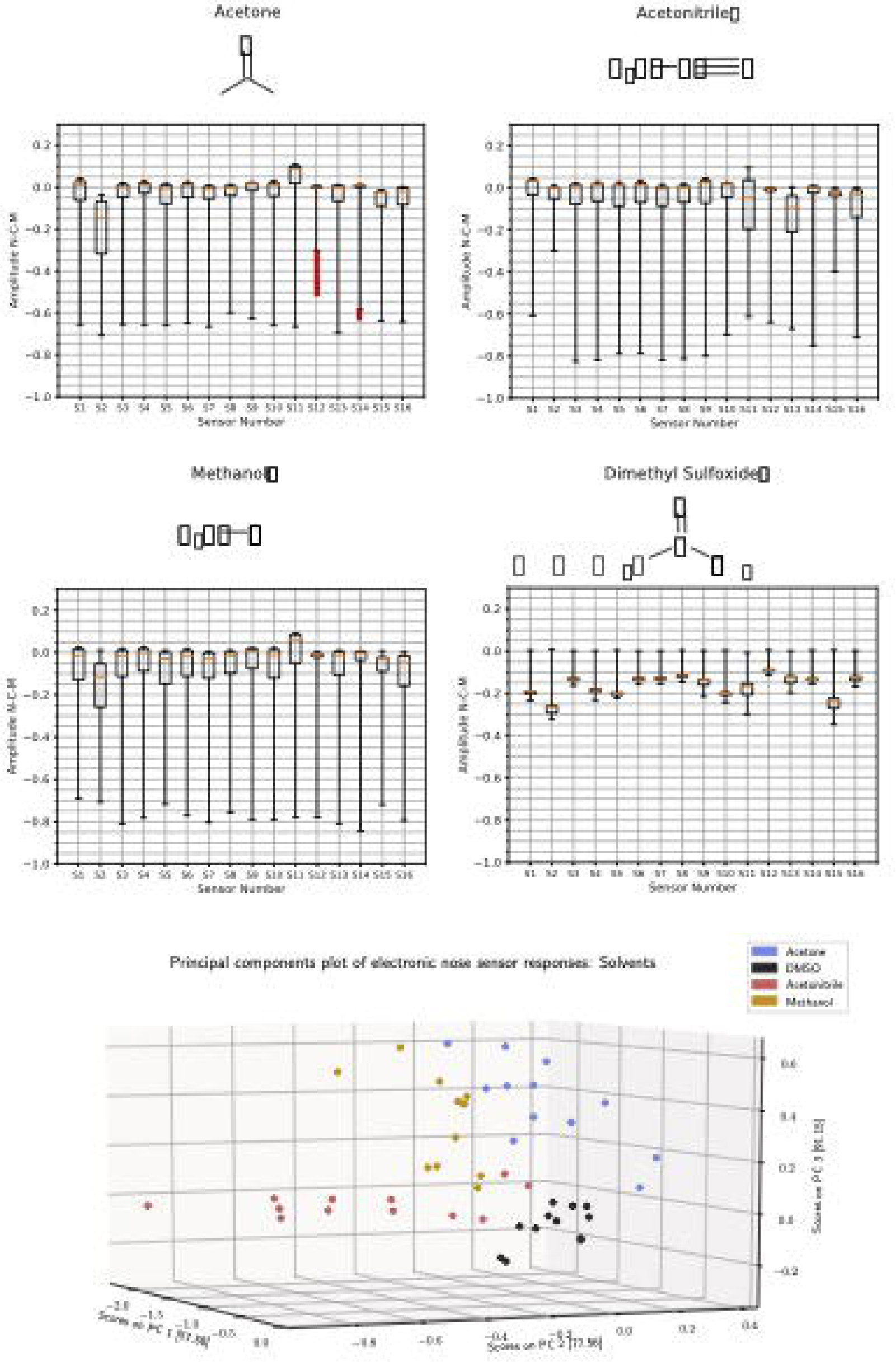
Electronic nose response with four different chemical solvents. Chemicals structures and olfactory response footprint which is the normalized centered and mean conductive amplitude of the sixteen sensors response: (**A**) acetone, (**B**) acetonitrile, (**C**) methanol, (**D**) dimethtyl-sulfoxide. (**E**) The first three principal components representation of the solvents.

The principal components analysis (PCA) obtained from the different tests with the chemical compounds (Fig. 1E) shows a sufficiently high sensorial response with the first three PCs (principal components), which correspond to 91.15% of the cumulative variance of the data. From this, a classifier based on vector support machines (SVM) based on the linear kernel was trained to quantify the separability in the response of each compound. The overall accuracy is 99.9%, an error of less than 1% to identify these compounds.

### Detection of Odor in the Presence of Triatomines

Most electronic nose prototypes are coupled to an independent pump cell as active airflow for better use of sensitivity and their properties. However, this model generates a degree of deoxidation in the detector material, generating a higher initial referential response of the gas sensors compared with the proposed model without a pump. Since an electronic nose with active independent airflow produces in the sensors a conductance response high enough to degrade sensitivity for detecting the low concentration of volatiles emitted by a single individual of triatomines, which is undesirable for our purposes. This observation, using an embedded active pump, was done as a preliminary evaluation as follows: Three adult non-fed females of the species *Triatoma dimidiata*, *Pastrongylus geniculatus*, and *Rhodnius prolixus* were used; the concentration-time, which means the time the incept remains in the container before applying the nose detection process, was 60 seconds and a container of 180 mL was used for saving the insects for the concentration-time (fig. S1A). As a result of this first experiment, low separability of the data was found using Principal component analysis (fig. S1B), so the inability of a nose, with an embedded active pump, to differentiate insect odors was demonstrated. That is why all the next experiments were done with a nose without an active pump.

The conditions of the following experiment to demonstrate the ability of the nose to detect a different kind of triatomines were as follows: The preparation of samples for experimental development was carried out with laboratory breeding colonies, whose population of triatomines was more remarkable than three individuals. The concentration time was more than three months, which is enough time for taking into account any amount of info-chemicals generated in a breeding habitat with different exuvial, corpses, eggs, and feces, among other natural processes of insects. The following kinds of genera and species of insects were used in the study: *Triatoma dimidiata, T. pallidepennis, T. maculata*, *Panstrongylus geniculatus, Rhodnius colombiensis*, *R. pallescens, R. prolixus* and *Eratyrus mucronatus*.

One treatment without insects was also analyzed to account for in the subsequent experiments. This means only the container alone was analyzed, including the cardboard base, fan-shaped cardboard, and ‘tulle veil’ or web. This is shown in Figure S3 (A to F) and explained in material and methods: subchapter ‘biologic material.’

Classificatory behavior was observed at the genus level and, in some cases, at the species level. Figure 2 (A to I) shows a representative image of each species followed by the smell footprint obtained, corresponding to the box plot of each sensor response in conductance, normalized, centered, and averaged.

**Fig. 2.**
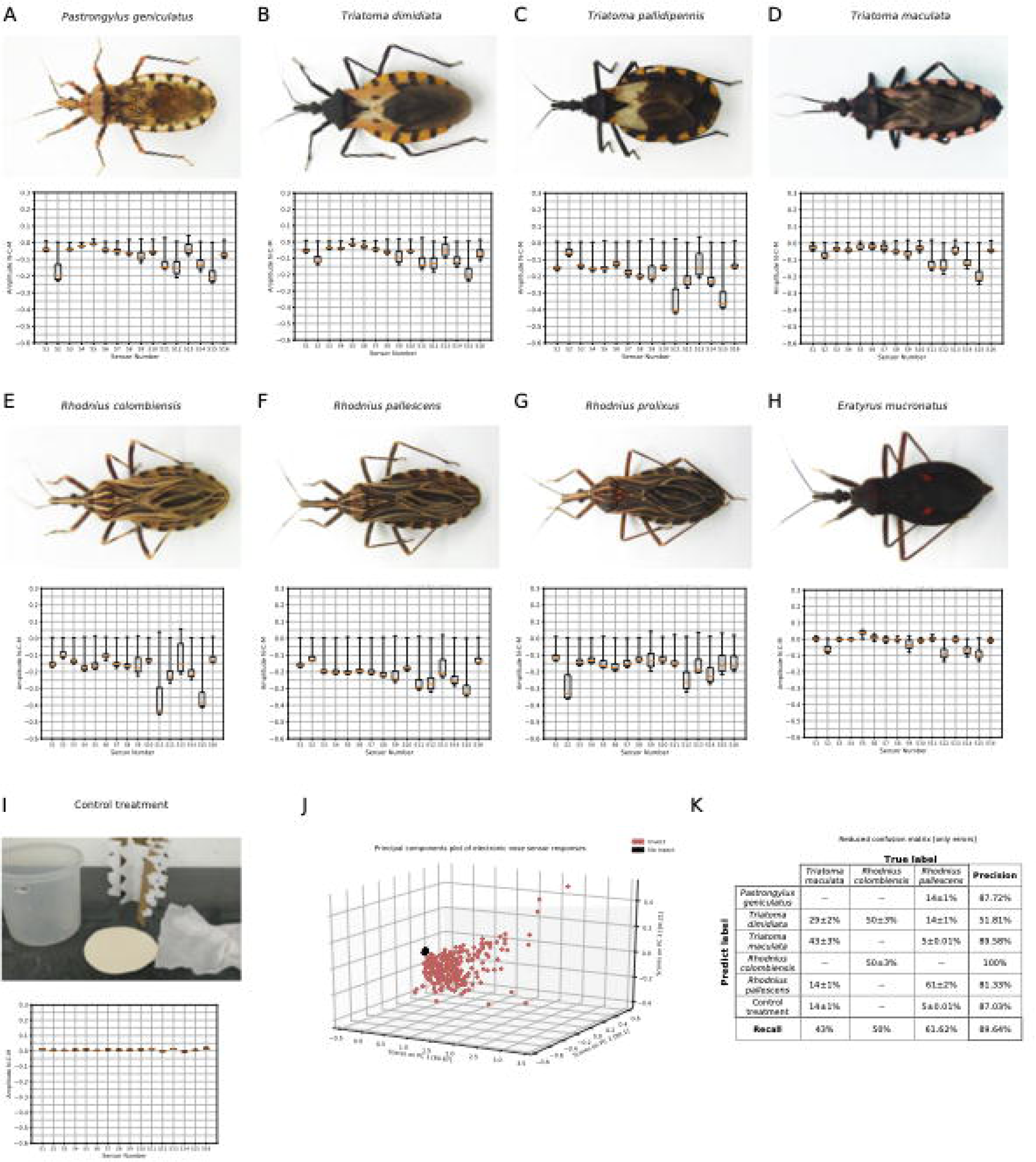
Electronic nose response with eight different species triatomines. Photographic representation and the normalized, centered and mean conductance amplitude response of eight triatominae species (**A**) *Pastrongylus geniculatus*, (**B**) *Triatoma dimidiata*, (**C**) *Triatoma pallidipennis*, (**D**) *Triatoma maculata*, (**E**) *Rhodnius colombiensis*, (**F**) *Rhodnius pallescens*, (**G**) *Rhodnius prolixus*, (**H**) *Eratyrus mucronatus*, (**I**) control treatment. (**J**) Representation of the first three principal components response that separate two groups: are or not insects (control treatment showed in the figure 2I and S3A). (**K**) Reduced confusion matrix table that shows only the labels with error occurred in the classification, it trained on the 8-class task, correct labels on the horizontal axis and detected label on the vertical axis.

During the experiments, a behavioral factor of the triatomines was observed before the caloric stimulus generated in the sensor chamber. This is because, in short distances (for this case, about 6cm), the thermal key (the heat emitted by a warm-blooded host) is a sufficient and necessary stimulus that allows the insects to orient themselves and trigger the harsh response 40, that is why in the figure S2A and movie S1 we can be appreciated the proboscis or triatomine horn through the web.

The principal components analysis (PCA) is plotted in three dimensions (the first three components) in Fig. 2J with a cumulative variance of 94.21%. A grouping behavior is observed in the data corresponding to the triatomine insect tests, separating from tests performed on elements without the presence of triatomine insects that can interfere with the structure of the colonies (fig. S3A) and the initial referential data of each sensor.

Figure 2K shows the confusion matrix of a multi-class support vector machine (SVM) with 9 output labels: eight triatomine species and one control treatment class; a general accuracy of 89.64% was obtained. In the case of the classification of samples of the species *T. maculata*, the classifier presented an error of 57% (recall 43%); that is, more than half of the samples were incorrectly classified, finding confusion between these species and *T. dimidiata* (29%), which are two species of the same genus. The accuracy of detection for *R. pallescens* is 81.33%, which indicates that there are outlier samples that share the same proportion with the species *T. dimidiata* (14%), *P. geniculatus* (14%) and in smaller proportion with *T. maculata* (5%) in addition to the control treatment (5%). On the other hand, the electronic nose classifies the genus Rhodnius with a precision of 79% because 38% of the samples of *R. pallescens* and 50% of *R. colombiensis* are poorly classified.

For the genus *Triatoma*, the electronic nose has a precision of 93.1%. However, the identification of the species *Triatoma dimidiata* presents a precision of 51.81% (also called positive predictive value: true positive / (true positive + true negative)) although the recall of prediction is more than 99% (also known as sensitivity: true positive / (true positive + false negative)). Finally, the samples were successfully classified with a maximum error lower than 0.1% for the cases where the data correspond to *Pastrongylus geniculatus*, *Triatoma dimidiata*, *Triatoma pallidipennis*, *Rhodnius prolixus*, *Eratyrus mucronatus,* and the control treatment.

### Variation in the Odor Concentration Time

Another important challenge to be faced was the need to explore the ability of the electronic nose to react to the volatiles emanated by the insects alone, as it is usually found in nature, without the biological waste that takes place in their environment during the concentration-time. Three main cases were tested: Case 1, where a three-month concentration time was used. The triatomines had to be saved in a container for three months before applying electronic nose detection. Case 2, was the same as case 1, but withdrew the insects away from the container. This means applying the test only to the biological waste (exuvia, corpses, feces, different impregnated odors, etc.). Case 3, where one-day concentration time was used, is equivalent to having insects free of any surrounding factors in the colony. In this case, the same insects of case 1 were used after a washing process on them was applied.

In other experiment conditions, the same genus and species were tested in each case. The three principal components of PCA plotting results for the three cases are shown in Figure 3 in this order: *P. geniculatus* (Fig. 3A), *T. dimidiata* (Fig. 3B), *T. pallidipennis* (Fig. 3C), *T. maculata* (Fig. 3D), *R. colombiensis* (Fig. 3E), *R. pallescens* (Fig. 3F), *R. prolixus* (Fig. 3G), *E. mucronatus* (Fig. 3H). The data processing corresponds to the conductance signals response of the 16 normalized, centered, and averaged nose sensors.

**Fig. 3.**
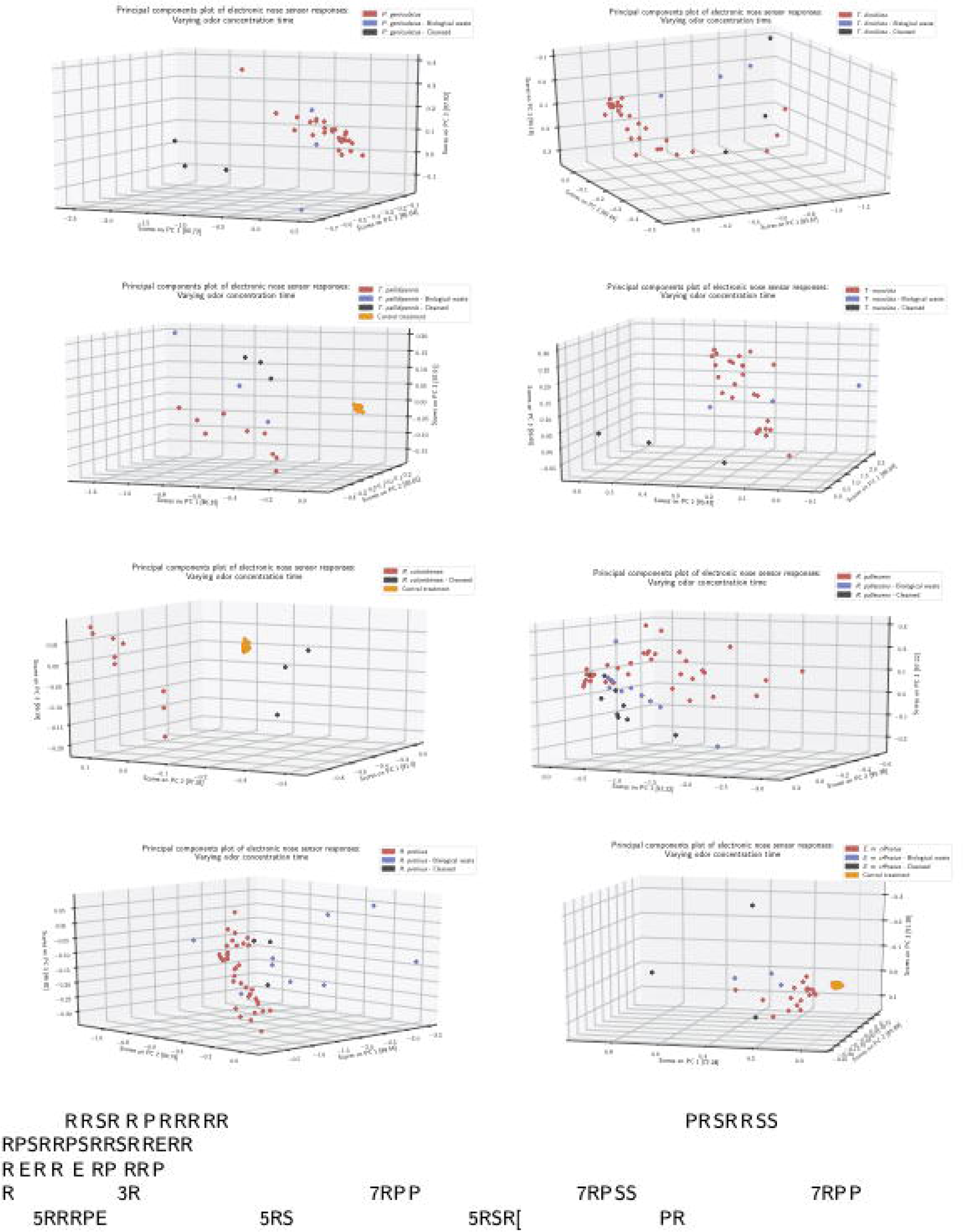
Electronic nose response with variation in time of odor concentration. A tridimensional plot of principal components analysis that compares the electronic nose response with individuals with waste (red), only biological waste without individuals (blue), only individuals without waste - cleansed (black) and in some cases the control treatment (yellow): (**A**) *Pastrongylus geniculatus*, (**B**) *Triatoma dimidiata*, (**C**) *Triatoma pallidipennis*, (**D**) *Triatoma maculata*, (**E**) *Rhodnius colombiensis*, (**F**) *Rhodnius pallescens*, (**G**) *Rhodnius prolixus*, (**H**) *Eratyrus mucronatus*.

The classification error between the colonies with more than three months of concentration and those with less than one day presents a high separability in most cases, as shown in Table 1. Specifically, this happened for the *P. geniculatus* (Tab. 1A), *T. dimidiata* (Tab. 1B), *T. pallidipennis* (Tab. 1C), *T. maculata* (Tab. 1D), *R. colombiensis* (Tab. 1E), *R. pallescens* (Tab. 1F), *R. prolixus* (Tab. 1G), and *E. mucronatus* (Tab. 1H). In case 2, the nose sensors’ behavior had the same tendency as in case 1, which means that the response in both cases corresponds mainly to the odors typical of waste and, to a lesser extent, to the odors directly emitted by the insects.

**Tab. 1.**
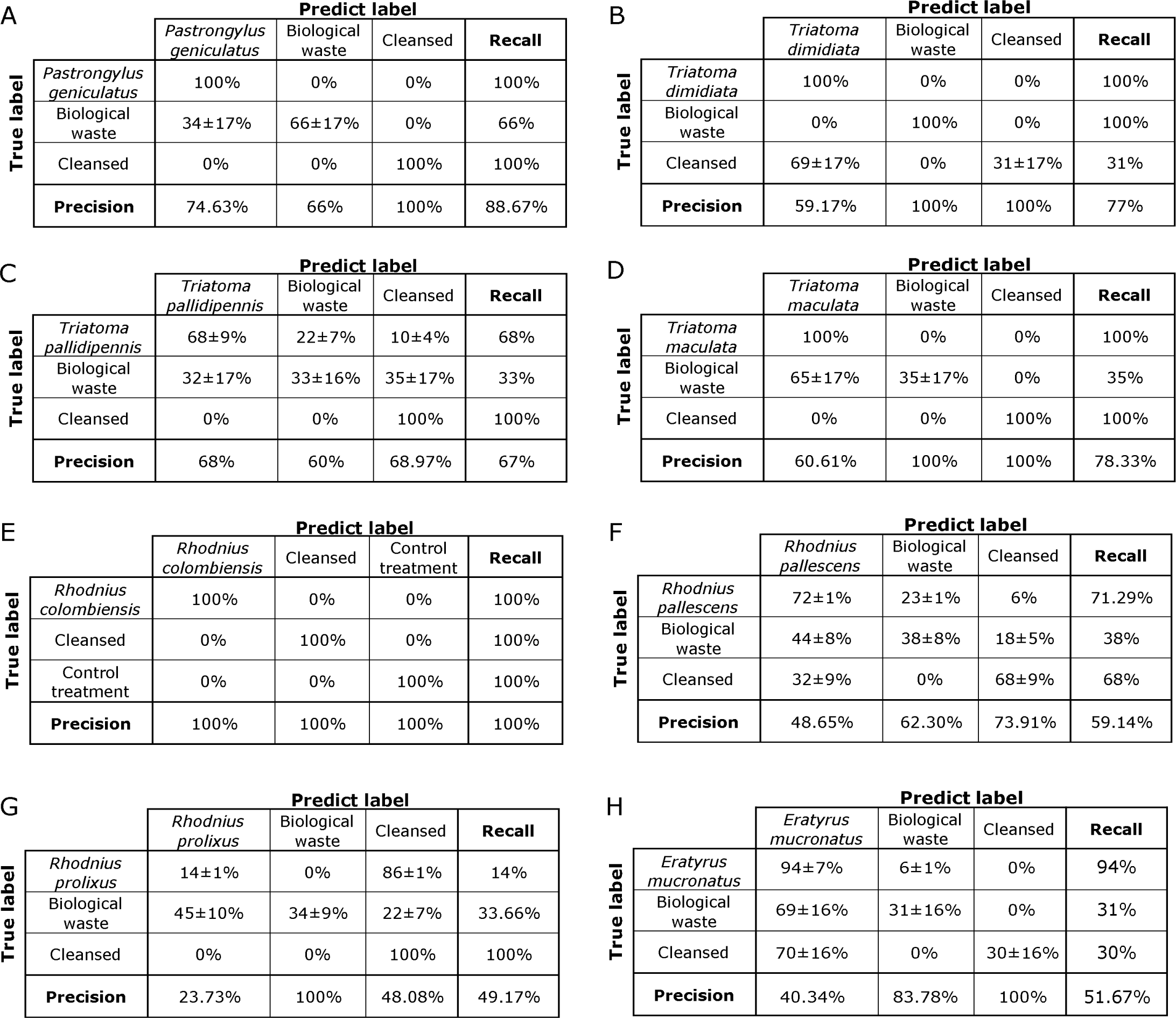
Confusion tables of electronic nose response with variation in time of odor concentration. Confusion matri table of sensors response trained on the 3-class task, the correct labels on the vertical axis and detected label on th horizontal axis. Comparasion of the electronic nose response with individuals and biological waste, only biologic waste without individuals, and only individuals without waste (cleansed): (**A**) *Pastrongylus geniculatus*, (**B**) *Triatom dimidiata*, (**C**) *Triatoma pallidipennis*, (**D**) *Triatoma maculata*, (**E**) *Rhodnius colombiensis*, (**F**) *Rhodnius pallescen* (**G**) *Rhodnius prolixus*, (**H**) *Eratyrus mucronatus*.

### Experimentation of Triatomine Specimens with the Red Eyes Genetic Variation

There are studies around triatomine species with several mutations in the literature. Among them, changes of color in the eyes have been reported that, according to authors, refer to an autosomal recessive mutation (Noé and Silva; Wygodzinsky and Briones; Vija-Suarez et al. 2017)41–43. Next, we present the results obtained from experiments using colonies from the triatomine species *Rhodnius prolixus* (Fig. 4A) and *Triatoma dimidiata* (Fig. 4B) with the red eyes genetic variation.

**Fig. 4.**
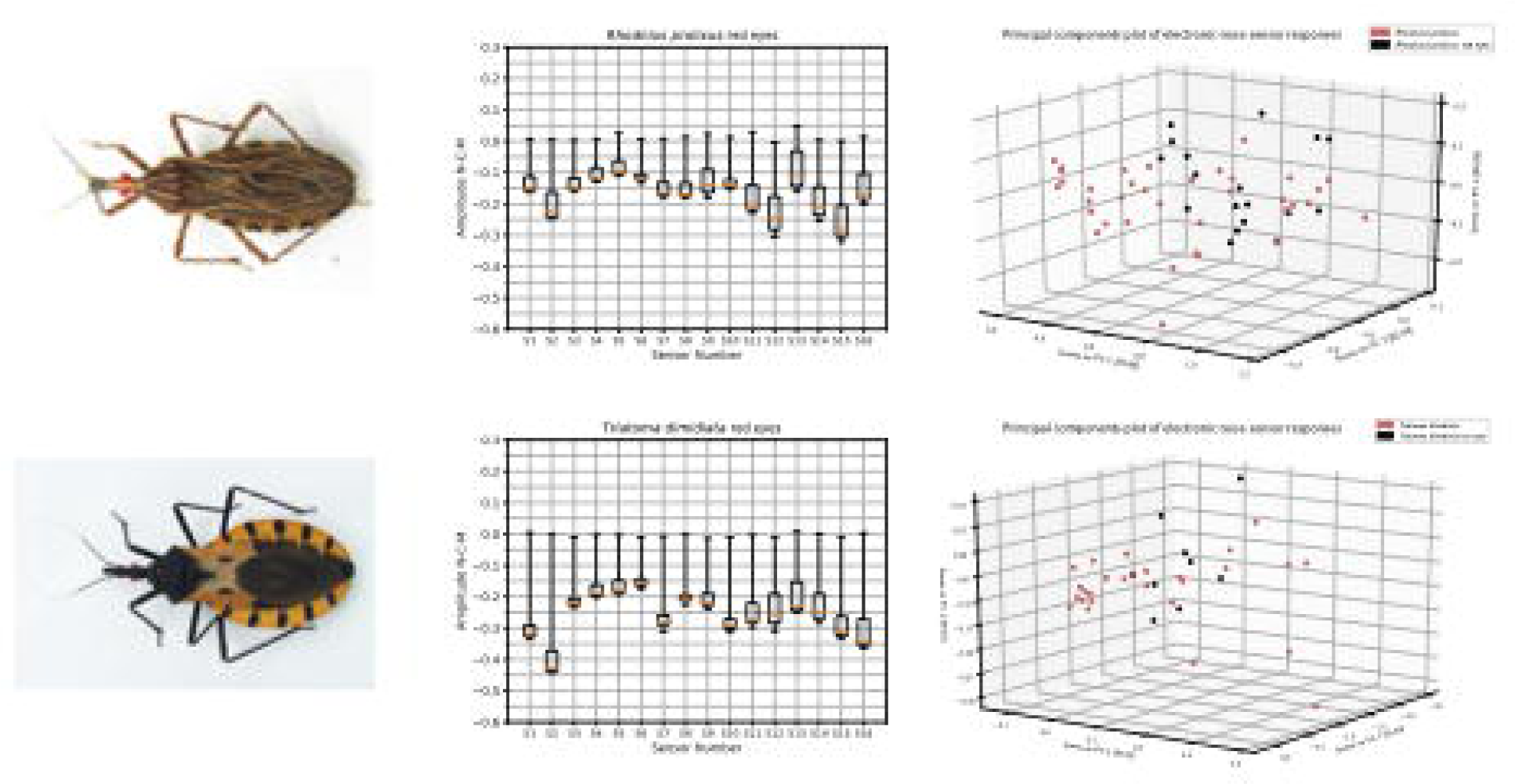
Experimentation of triatomine specimens with the red eyes genetic variation. Photographic representation of (**A**) *Rhodnius prolixus* red eyes and (**B**) *Triatoma dimidiata* red eyes followed by footprint odor of (**C**) *Rhodnius prolixus red eyes* and (**D**) *Triatoma dimidiata* red eyes. The analysis of the principal components of (**E**) *Rhodnius prolixus* red eyes and (**F**) *Triatoma dimidiata* red eyes.

In agreement with the previous experiments, Figure 4 (A and B) shows the photographs of two individuals from each colony used in the experimentation as visual representatives. In Figure 4 (C and D), the smelling trace obtained is shown, which corresponds to the conductance response signals of the 16 normalized, centered, and averaged sensors. There is a low separability between the trials belonging to each of the two red eyes’ genetic variation species compared to their referents without the mutation. This is evidenced by the principal components analysis (PCA) with the first three PCs for both the *R. prolixus* (Fig. 4E) and the *T. dimidiata* (Fig. 4F) species. In the first case, the PCs accumulate 88.09% and 96.82% of the total data variance for the second.

The behavior of the support vector machine (SVM) with linear kernel presented an accuracy to differentiate between the tests carried out on species *R. prolixus* with genetic variation and without the genetic variation of 69.3%. In the cases of *T. dimidiata* trials, the accuracy corresponds to 58.1%, that is, a low separability in the data. There was no confusion between species *R. prolixus* and *T. dimidiata* (with the genetic variation); in this case, the error is less than 0.1%, and the results are shown in the section “Detection of odor in the presence of triatomines.”

### Differentiation in the Smell of Triatomines, Mosquitoes, and Rice Weevils

In the case of the treatments related to the species *A. aegypti*, a typical behavior of the mosquitoes was observed. Eventually, they were flying and posing on the walls and the container’s web, as shown in Figure S2B. The oxidation behavior of SnO2 material in 12 of the 16 sensors was observed in the obtained data; that is, an increase in the resistance is a particular behavior of this treatment.

In the case of the *S. oryzae* colonies, during the experimentation, aggregation behavior was observed on the edges of the container just after finishing the acquisition of data. Some individuals abandoned the rice in the bottom and moved towards the source of heat (sensors) trapped in the web. See figure of supplementary materials figure S2C.

Figure 5A shows an individual representative of the *A. aegypti* colony, and Figure 5B shows an individual of *S. oryzae*. In Figure 5 (C and D), we can see its odor trace followed by the three-dimensional graph of the analysis of principal components (Fig. 5E) of the tests carried out with mosquitoes, weevils, and the eight species of triatomine shown above in the section “Detection of odor in the presence of triatomines.”

**Fig. 5.**
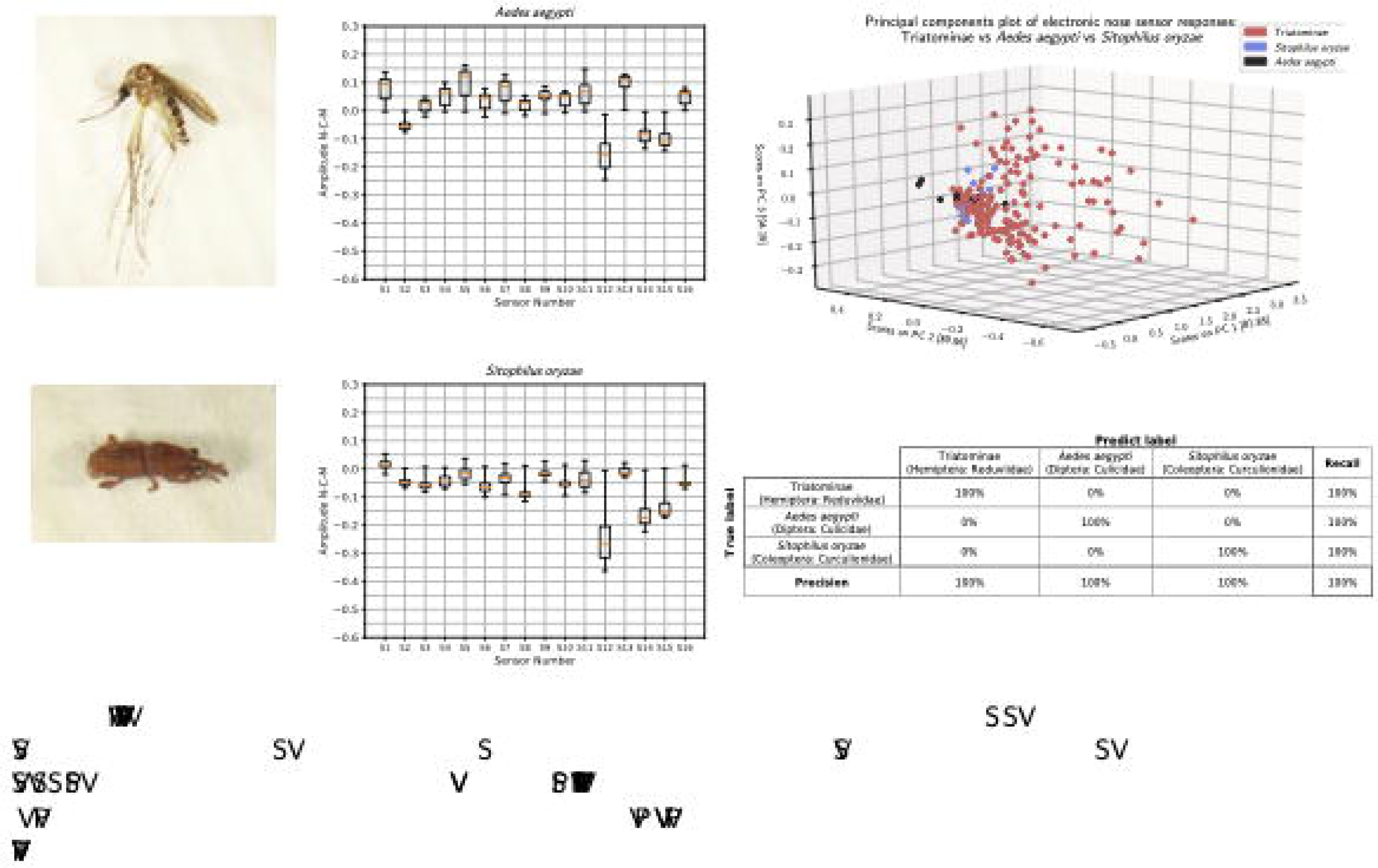
Differentiation in the smell of triatomines, mosquitoes and rice weevils. Photographic representation of (**A**) *Aedes aegypti* and (**B**) *Sitophilus oryzae* followed by footprint odor (**C**) *Aedes aegypti* and (**D**) *Sitophilus oryzae*. (**E**) The response of the principal components analysis that compare the smell between triatomines, mosquitoes and weevils, can be observed in the three-dimensional drawing. (**F**) The confusion matrix table of the classifier between triatomines, mosquitoes, and weevils

Although it is impossible to show high separability in the response with the first three principal components, which represent 94.19% of the cumulative variance of the original data, 10 PCs were used for the classification. Finally, the classifier based on the vectorial support machine succeeds in identifying and separating the colonies belonging to the subfamily of Triatominae from the colonies belonging to the species *A. aegypti* and *S. oryzae* with an error of less than 1% (Fig. 5F). In the same way, all the samples that correspond to the control treatment were correctly classified.

### Differentiation of Colonies Infected with the *Trypanosoma Cruzi* Parasite

It is of great interest to identify the insect vector of Chagas disease and know if the triatomine is infected with the parasite *Trypanosoma cruzi*. For this reason, tests were conducted with the electronic nose for the species *Rhodnius pallescens* which had been naturally infected. The detection of *T. cruzi* was performed by amplification by PCR of a partial region of the heat-shock protein 70 gene (hsp70) of *T. cruzi* using specific primers and PCR conditions previously reported, as depicted in Figure 6A.

**Figure 6.**
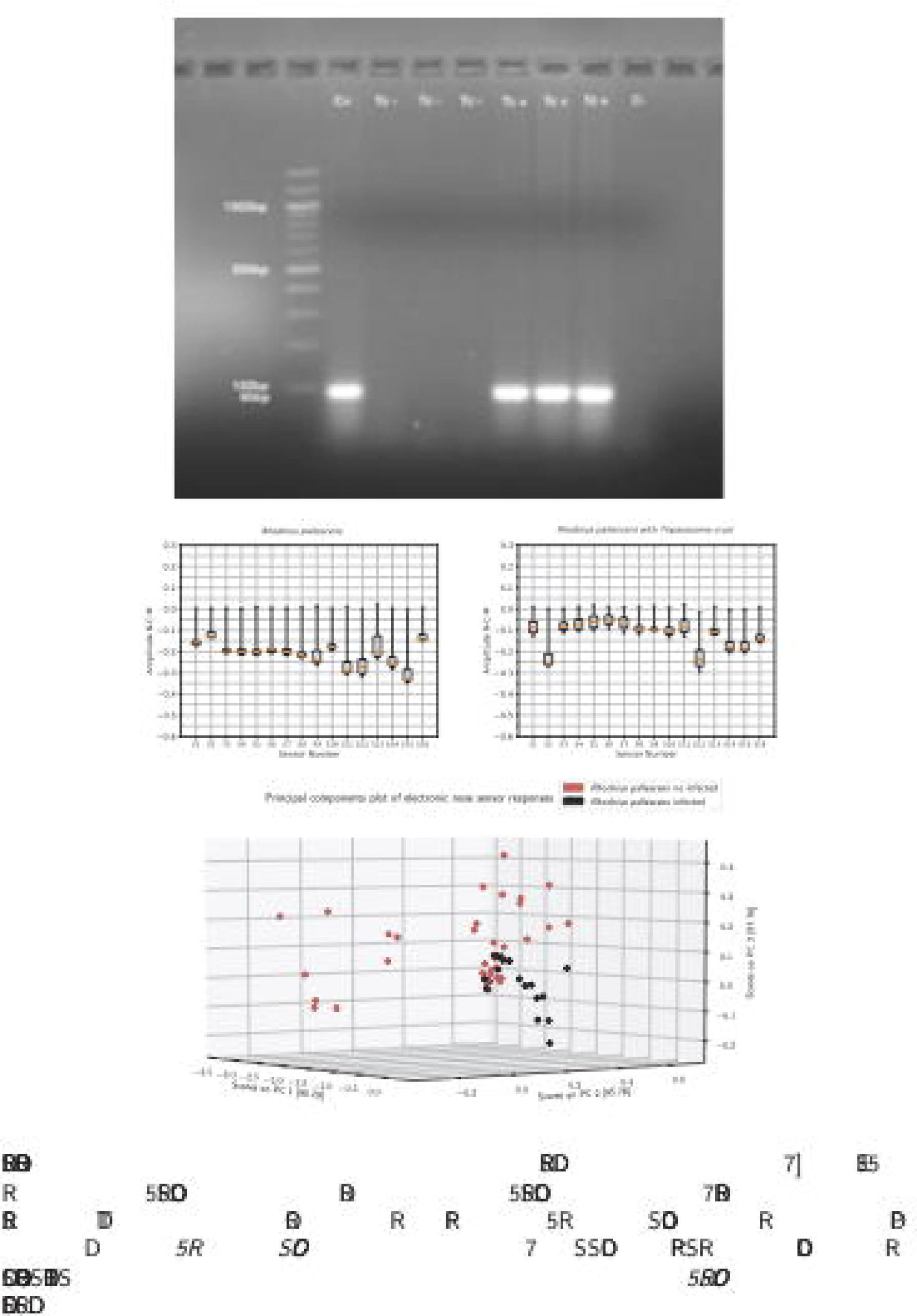
Differentiation of infected colonies. Detection of the parasite *T. cruzi* by PCR in feces of (**A**) *Rhodnius pallescens* non infected and *R. pallescens* infected. The odorant footprint acquired with the electronic nose from (B) *R. pallescens* non-infected (same in Fig.2F) and (**C**) *R. pallescens* infected. (**D**) The principal components analysis showing relative separability between volatile samples acquired from the species *R. pallescens* with absence ad presence of the parasite.

In addition, in Figure 6B, the odoriferous trace obtained from the colonies without the parasite can be observed, and in Figure 6C, with the test colonies where the *T. cruzi* parasite was found in the feces of the triatomines. In the analysis of principal components shown in Figure 6D, separability between the acquired data is evidenced that insects are parasitized. In this way, the classifier’s training indicated an error of less than 1% in the classification. In the same way, all the samples that correspond to the control treatment were correctly classified.

## Discussion

An electronic nose version is presented under the IoT (Internet of Things) platform mobile terminal device coupled with artificial intelligence in the cloud, designed to be a versatile, portable, trainable, low-cost device with access to multiple users (Gardner and Bartlett 2013). The electronic nose as IoT devices facilitate the task of using them to identify possible insects carrying Chagas disease. Some qualities of the device presented in this investigation are: (i) It does not affect the experimental individuals, (ii) The risks when handling insects either in the training phase or in the detection phase of the electronic nose are reduced. (iii) The device is comfortable compared to the commonly found electronic noses due to its portability. (iv) Nose detection results are obtained immediately and economically compared to the morphological identification of the triatomines, which is slow because of the need for an expert in the phylogeny of the animal to analyze the morphological characters. (v) The time the training was generated was short compared with other methodologies. On the other hand, Its training is constant, which means every time a new system feature is sniffed out, it will learn it. (vi) The hardware is simple, buildable, replicable, and modifiable since it is built-in sequential modules and a specific design can be made according to the working conditions.

The described work emphasizes the technological importance of developing complementary senses for humanoid robots. One is the olfactory sense, which has future applications in public health, such as the possibility of vector detection/identification. In this case, an electronic olfactory system was evaluated to detect and differentiate smells expelled by insects of medical importance, like the need to identify the Chagas disease vector.

Current research on humanoid robotics is fused with artificial intelligence, which involves the capability of the robots to respond to the natural dynamic change in their environment (Ganesan et al. 2006; Yeon et al. 2015; Wang et al. 2015). This area covers significant advances in replicating vision (image processing and object recognition), hearing, touch, taste (using electronic tongues), smell, communication ability, and many other aspects currently associated with humans. As a result, new opportunities are opened, like the possibility of working in the future in the field of prevention, surveillance, and control of contagious diseases with the help of artificial intelligence.

Our device presents a more straightforward solution compared with other common electronic noses (Moreno et al. 2009), in terms of modularity and technical specifications that allow it to adapt to a wide range of robotic devices. The three types of interactions have been pointed out: chemistry, electronics, and logic support (software and learning algorithms), so our nose has been optimized in its cost, size, portability, and effectiveness using the advances in sensor technology, electronics, programming, and cloud computing. Consequently, a future swarm of internet-connected electronic noses is also possible for many other purposes.

On the other hand, the designed electronic nose allowed us to detect the presence or absence of triatomines without specifying information about the chemical compounds in the environment where insects live. Nevertheless, the most important fact is that each kind of triatomine has a clear-smelling footprint that makes it recognizable, as demonstrated throughout this research.

On the other hand, using an electronic nose requires a special methodology. The first phase is the validation phase of the operation of the electronic nose. The purpose is to test the reaction of the sensor to some specific solvents and take the results of this as a performance indicator and as a reference point for future procedures. Their selection of the solvents was based on their properties to react with the material of each of the sensors. Some of these properties are the polarity, the number of free electrons capable of creating bonds with oxygen present in the material of a sensor, and also the boiling point of a solvent, which is responsible for its volatility, which is important. Following this precedence, the solvents proposed to be used are, in order of their volatility, acetone (56 ° C) followed by methanol (65 ° C), acetonitrile (82 ° C), and finally the dimethyl-sulfoxide or DMSO (189 ° C).

The volatility of the four solvents is evidenced in the results shown in Figure 1(A-D). Figure 1A belongs to the case of the use of acetone. A clear footprint is seen in the Figure for this solvent as follows: For the sensor S2 (tagged as MQ3), it takes too much more time than others to reach the higher level when the solvent is present, which appears as a large box in the Figure; It takes also too much more time than others to recover to the basal state (cero level), when the solvent is evaporated, which is shown in Figure 1A, as a box located far from the cero level; Sensors S12 (tagged TGS-816) and S14 (tagged TGS-813) behaves as outliers, because their recovery speed, to the basal state, is fast enough to appear as the red vertical line in Figure 1A; The other sensors has a regular behavior with an intermediate time to react and to recovery to the basal state. The behavior of sensor S2 is due to the doping of the material, which makes it highly sensitive to gasses like methane, LPG, smoke, alcohol, ethanol, hexane, and CO, and indicates an area of common use oriented to the detection of alcohols and solvents.

The other cases will be explained by comparing the use of acetone, acetonitrile, methanol, and DMSO, see Figure 1(A-D). Each footprint has its range of amplitude which is the difference between the maximum and the minimum value of the amplitude of the sensor signal. In the case of acetone, acetonitrile, and methanol, the amplitude range is higher than that obtained by DMSO. The separability of these compounds is high (error less than 0.1%), indicating that the electronic nose has no difficulty differentiating all these solvents. Similar results of the investigation can be found in the literature (Rodriguez-Lujan et al. 2014; Sanz Montero 2019; Abeywickrama et al. 2023).

Experiments with specimens of the Triatominae subfamily indicated that the odor generated during a long concentration time (more than three months) is different from the smell of an environment free from the presence of the insect, limited to laboratory conditions (shown in materials and methods). This indicates the ability of the electronic nose to react differently to different environments.

The measurements among the eight species used yielded high levels of separation not only by gender but, in some cases, also by species. According to some studies, the chemical composition of the volatiles, mainly of the exocrine glands of different triatomines, presents isobutyric acid and, according to other studies, acetic acid as a major component (Rojas et al. 2002; Guerenstein and Guerin 2004; May-Concha et al. 2015). However, it has been shown that, in general, a large mixture is formed of a short chain of fatty acids such as esters, and alcohols, and components such as ketones (Minoli et al. 2013). It is well known that the sexual aspect is one of the taxonomically classificatory characteristics. Therefore it would be expected that the volatile chemical substances associated with the reproduction of pheromones of each species will be different even though they present the same majority of compounds (Pattenden and Staddon 1972; Games et al. 1974; Kälin and Barrett 1975; Guarneri and Lorenzo 2021). In addition, nutritional status is also an essential variable in the secretion of volatiles, as well as aspects such as the correlated determination of the population density and the concentration time of the odors. These factors can respond to the classification errors in *R. pallescens*, *R. colombiensis*, and *T. maculata* samples.

Reducing the concentration time from three months to less than a day was necessary to verify the effects of the natural interferences associated with the info chemicals developed in the test colonies, thus separating the biological waste from the live insects. Chemical signs of aggregation have been found in triatomine feces (Schofield and Patterson 1977; Pontes et al. 2008; Vitta et al. 2009), which allows insects to locate hosts and recognize whether or not they are safe. During oviposition, the females secrete volatiles to indicate the existence of the place where the eggs were deposited (Hubbard et al. 1987; Gürtler et al. 1999; Ganesan et al. 2006). These additional volatiles, together with the unique presence of the triatomine insects in the colonies, are evidenced when performing the experimentation of separating the biological wastes. It was found that the nose can detect the waste of the insects rather than the insects themselves.

We found that in the two cases studied with red eyes genetic variation, *Rhodnius prolixus,* and *Triatoma dimidiata*, no volatiles are generated that react with the materials of the sensors in a different way to the results obtained with the colonies without the genetic variation. It has been reported that this mutation is autosomal recessive (Wygodzinsky and Briones; Vija-Suarez et al. 2017), and there is no evidence in the literature of metabolic changes in insects that may indicate the generation of particular and measurable chemical compounds through the sensors implemented in the proposed prototype.

On the other hand, the ability of the electronic nose to find differences in volatiles emanating from insects of another taxonomic order, which belong to Coleoptera and Diptera, was verified. The morphological characteristics, the alimentary and sexual habits, among other characteristics, were indirectly measured by the electronic olfaction system, thus differentiating colonies of the species *Aedes aegypti (Ganesan et al. 2006; Chotiwan et al. 2018)* and *Sitophilus oryzae* (Phillips et al. 1987; Collins et al. 2007) from the triatomines, with a low error of 0.1%.

In the same way, the electronic nose found differences between colonies of *R. pallescens* infected and not infected. Hence, the triatomines with the presence of the parasite *Trypanosoma cruzi* have a clear footprint that makes them distinguishable from others. Studies conducted with different species of triatomines have shown that the parasite generates metabolic changes in the insect (IEEE Staff 2010; Zumaya-Estrada et al. 2018; Erny and Masuda 2022), due to interactions between the insect intestine and the parasite in its different life stages.

Finally, the measurements of odors that allow the detection of the vectors of Chagas disease, with approximations to the identification of the presence of the parasite in transmitting insects, provide valuable information coupled to a humanoid robot. In the future, this setting will allow for locating the source of the odor and therefore identify infected insects. From this perspective, our next phase of work will focus on scaling the electronic nose in conditions outside the laboratory, see Figure 7, closer to practice, evaluating the possible environmental and natural smell scents where a colony of triatomine insects is possible. On the other hand, future work will also focus on a system of multiple electronic noses as vector detectors while emphasizing communication between swarm-type noses.

**Fig. 7.**
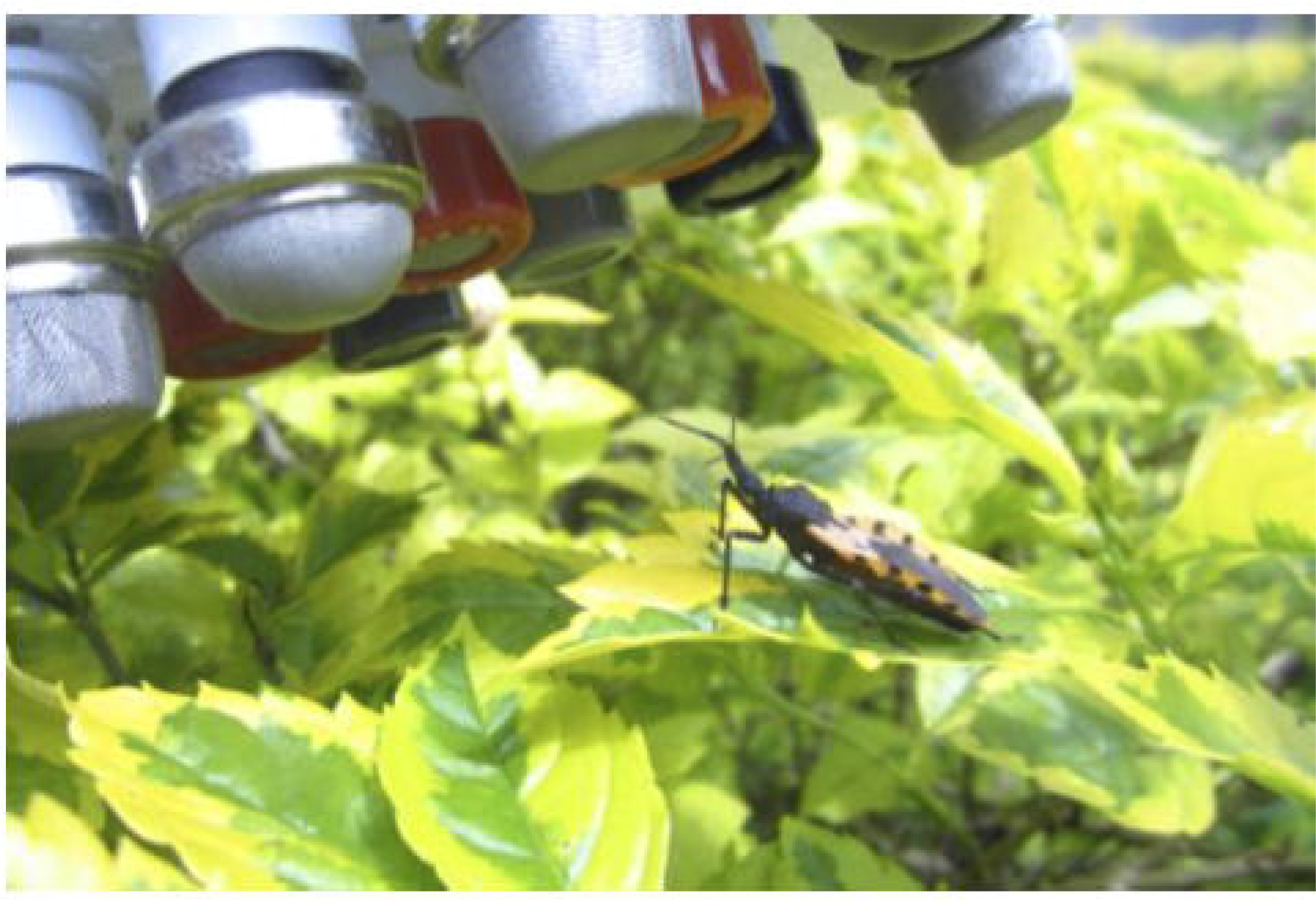
The electronic nose for triatomine detection in natural area. Future work representative image

## Acknowledgements

We thank the members of the Tropical Diseases Research Center (CINTROP) in the Universidad Industrial de Santander (Colombia), in particular, G. Rincon, M. Vidal, and R. Castillo. We thank all who were involved in the development of the electronic nose and the treatment of the biological material. We thank Prof. Oscar Eduardo Gualdrón from Universidad de Pamplona (Colombia) for his support and advice in developing the electronic nose.

## Author contributions

L.F.R. developed the hardware for the electronic nose, collected and processed the data, and drafted the manuscript. D.A.S designed and supervised the data processing. H.O.B. assisted in implementing the hardware and communication protocol. B.R.O. helped in the detection of *T. cruzi* by PCR. J.E.D. investigated the proposed approach and supervised the research. All authors revised and contributed to the manuscript.

## Data and materials availability

All data needed to support the conclusions of this research can be found in the manuscript, and supplementary materials are available online as a supplementary file S6. Contact J.E.D for more materials.

